# Antibiotic-induced microbiota disruption impairs neutrophil-mediated immunity to respiratory *Aspergillus fumigatus* infection in mice

**DOI:** 10.1101/2025.11.18.689104

**Authors:** Mariano A. Aufiero, Tobias M. Hohl

## Abstract

*Aspergillus fumigatus* is an environmental mold that forms ubiquitous airborne conidia and can cause life-threatening infections in immunocompromised individuals. Invasive aspergillosis occurs in patients with quantitative or qualitative neutrophil defects, who often receive systemic antibiotics to prevent or manage bacterial infections. Antibiotic-induced bacterial dysbiosis has been linked to impaired neutrophil bactericidal activity and to intestinal commensal bacteria escape during systemic candidiasis, though it remains unclear whether receipt of antibacterial antibiotics impairs neutrophil-dependent defenses against inhaled mold pathogens in the lung. Herein, we measured the outcome of *Aspergillus* challenge in C57BL/6J mice that were treated with different antibiotics in the drinking water for three weeks prior to experimental infection. We found that ampicillin but not neomycin or vancomycin treatment significantly increased murine mortality and lung fungal burden. The heightened susceptibility was associated with impaired fungal killing by lung-infiltrating neutrophils and monocytes, as well as reduced neutrophil production of NADPH oxidase 2 (NOX2)-dependent reactive oxygen species (ROS). These findings demonstrate that systemic antibiotic treatment can compromise pulmonary anti-*Aspergillus* immunity and suggest that the host microbiota can enhance neutrophil fungicidal activity by promoting NOX2-mediated ROS production.

**IMPORTANCE:** *Aspergillus fumigatus* is an environmental mold that causes invasive pulmonary disease in immunocompromised individuals. Owing to limited diagnostic tools, a narrow arsenal of effective treatments, and rising antifungal resistance, the World Health Organization (WHO) has designated *Aspergillus* as a critical priority fungal pathogen, highlighting the urgent need for further research. Patients with compromised immunity often receive broad-spectrum antibiotics to prevent or treat opportunistic infections, leading to significant disruption of the resident commensal microbiota. This antibiotic-induced dysbiosis has been linked to *Clostridium difficile* colitis and to intestinal overgrowth of vancomycin-resistant *Enterococcus* and *Candida parapsilosis*, preceding bloodstream infection. However, the impact of antibiotic treatment on susceptibility to invasive pulmonary aspergillosis remains undefined. In this study, we found that oral treatment with ampicillin, but not neomycin or vancomycin, significantly increased mortality in mice following *Aspergillus* infection. Neutrophils from the lungs of ampicillin-treated mice also showed markedly impaired fungal killing. These findings raise the possibility that preserving microbiome integrity during antibiotic treatment could enhance immune protection against invasive aspergillosis in at-risk patient groups.

## OBSERVATION

Phagocytes such as neutrophils, inflammatory monocytes, and alveolar macrophages are key mediators of host defense against *Aspergillus* (1–3). Studies in murine models of *S. pneumoniae* and influenza virus infections have demonstrated a link between the composition of the intestinal microbiota and phagocyte function during acute pulmonary infections (4, 5). Additionally, antibiotic treatment results in susceptibility to bloodstream and oropharyngeal infection with *Candida albicans* (6, 7). These studies demonstrate that antibiotics shape pulmonary phagocyte responses and influence susceptibility to fungal infection. However, the impact of antibiotic treatment on host defense during *Aspergillus* infection and how this affects disease progression remain unclear.

To test if antibiotic treatment increases susceptibility to *Aspergillus* infection, we treated C57BL/6J (WT) mice with one of three distinct classes of antibiotics in the drinking water for 3 weeks. Antibiotics were withdrawn 1 day before intratracheal (i.t.) infection with conidia (Figure 1A). As a control, WT mice were housed in our colony but received normal drinking water (designated NT). Vancomycin, a glycopeptide which primarily targets Gram-positive bacteria, or neomycin, an aminoglycoside which primarily targets Gram-negative bacteria, had no effect on survival following pulmonary *Aspergillus* infection (Figure 1B). However, treatment with ampicillin, a β-lactam antibiotic that targets Gram-positive, Gram-negative and anaerobic bacteria that lack β-lactamase activity, resulted in increased mortality following *Aspergillus* challenge (Figure 1C). To assess whether this susceptibility reflected a higher organ fungal burden, we measured *Aspergillus* lung CFUs 36 hours post infection (hpi) and found an increase in ampicillin-treated but not in vancomycin- or neomycin-treated mice (Figure 1D). Histopathology analysis revealed fungal tissue invasion in ampicillin-treated lungs (Figure 1E). Together, these findings indicate that ampicillin treatment increases mortality due to increased lung fungal burden.

**Figure 1:**
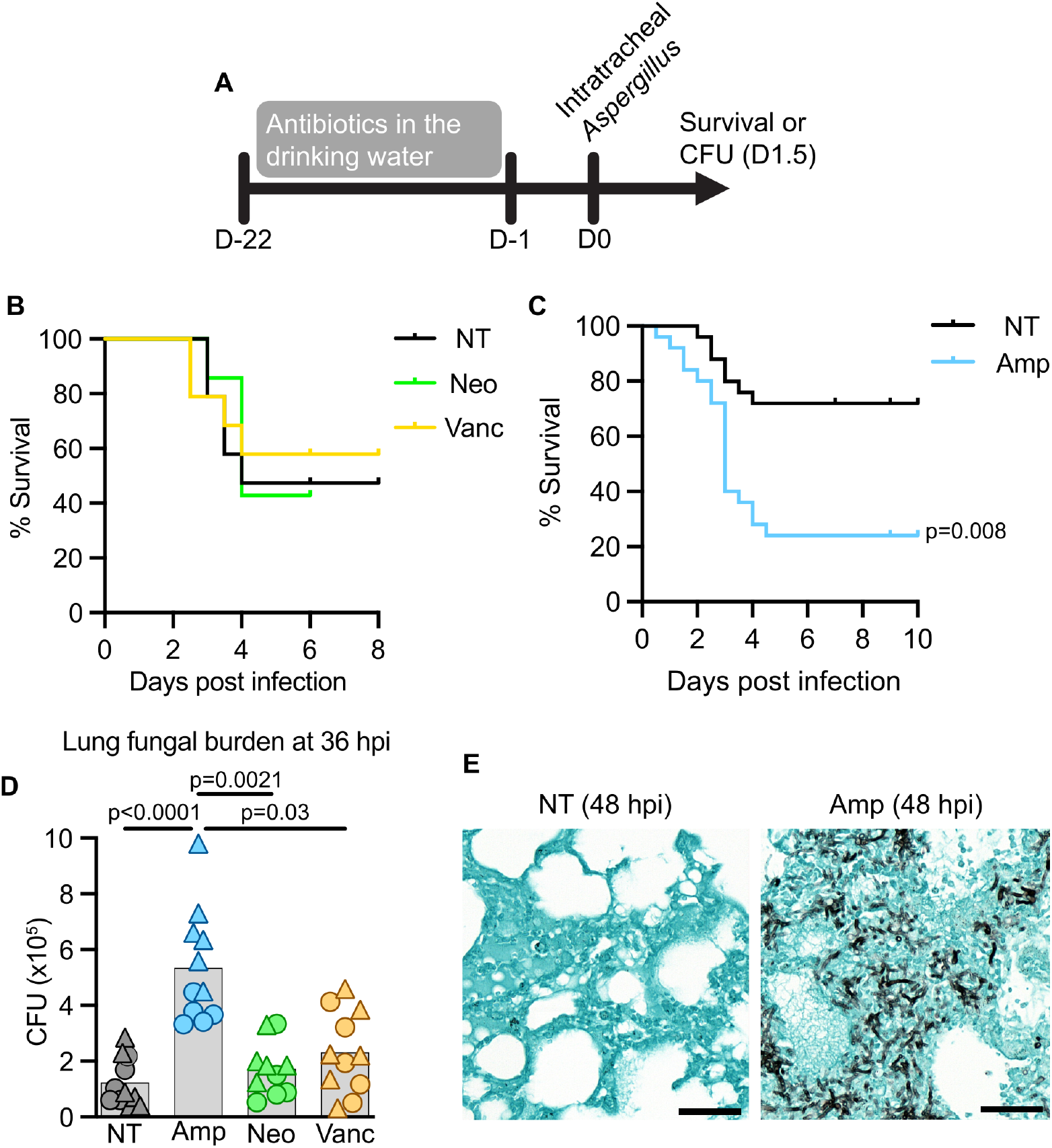
Antibiotic treatment increases susceptibility to *Aspergillus* infection. (A) Schematic of the experimental timeline. (B and C) Murine survival following i.t. infection with *Aspergillus* conidia, comparing not-treated (NT) mice to mice treated with neomycin and vancomycin (B) or ampicillin (C). Data are pooled from at least 2 experiments. n = 17 in (B) and n = 25 in (C). (D) Fungal burden in the lungs at 36 hours post infection, as quantified by CFUs. Data are pooled from two experiments and point shape indicates experiment of origin. (E) Representative histology (Grocott–Gömöri’s methenamine silver stain) of lungs from control and ampicillin-treated mice 48 hours post infection. Statistics: (B and C) Log-rank (Mantel–Cox) test; (D) Kruskal–Wallis test with Dunn’s multiple comparisons test.

Fungal growth in the lung often reflects defective recruitment or function of neutrophils or inflammatory monocytes (2, 7). To test this, we quantified these cells in the lungs of untreated and ampicillin-treated mice after infection and found no difference at 36 hpi (Figure S1A and S1B). We then used Fluorescent *Aspergillus* Reporter (FLARE) conidia to assess fungal uptake and killing by lung neutrophils and monocytes, as previously described (8). FLARE conidia encode mRFP and are uniformly labeled with Alexa Fluor 633 (AF633) as a tracer fluorophore. FLARE conidia shift their fluorescence emission from AF633^+^mRFP^+^ to AF633^+^mRFP^−^ when conidia are inactivated within lung phagocytes. Neutrophil and monocyte conidial uptake, i.e. the frequency of neutrophils and monocytes that contain either AF633^+^mRFP^+^ to AF633^+^mRFP^−^ conidia, was not significantly different between antibiotic-treated and untreated mice (Figure 2A– 2C). However, lung neutrophils and monocytes from ampicillin-treated mice contained a higher fraction of mRFP^+^ conidia than those from untreated or other antibiotic-treated mice (Figures 2D–E), indicating a fungal killing defect. Thus, lung fungal clearance in ampicillin-treated mice is impaired due to defective myeloid cell fungal killing, but not defective myeloid cell uptake nor defective myeloid cell recruitment and/or survival in the lung.

**Figure 2:**
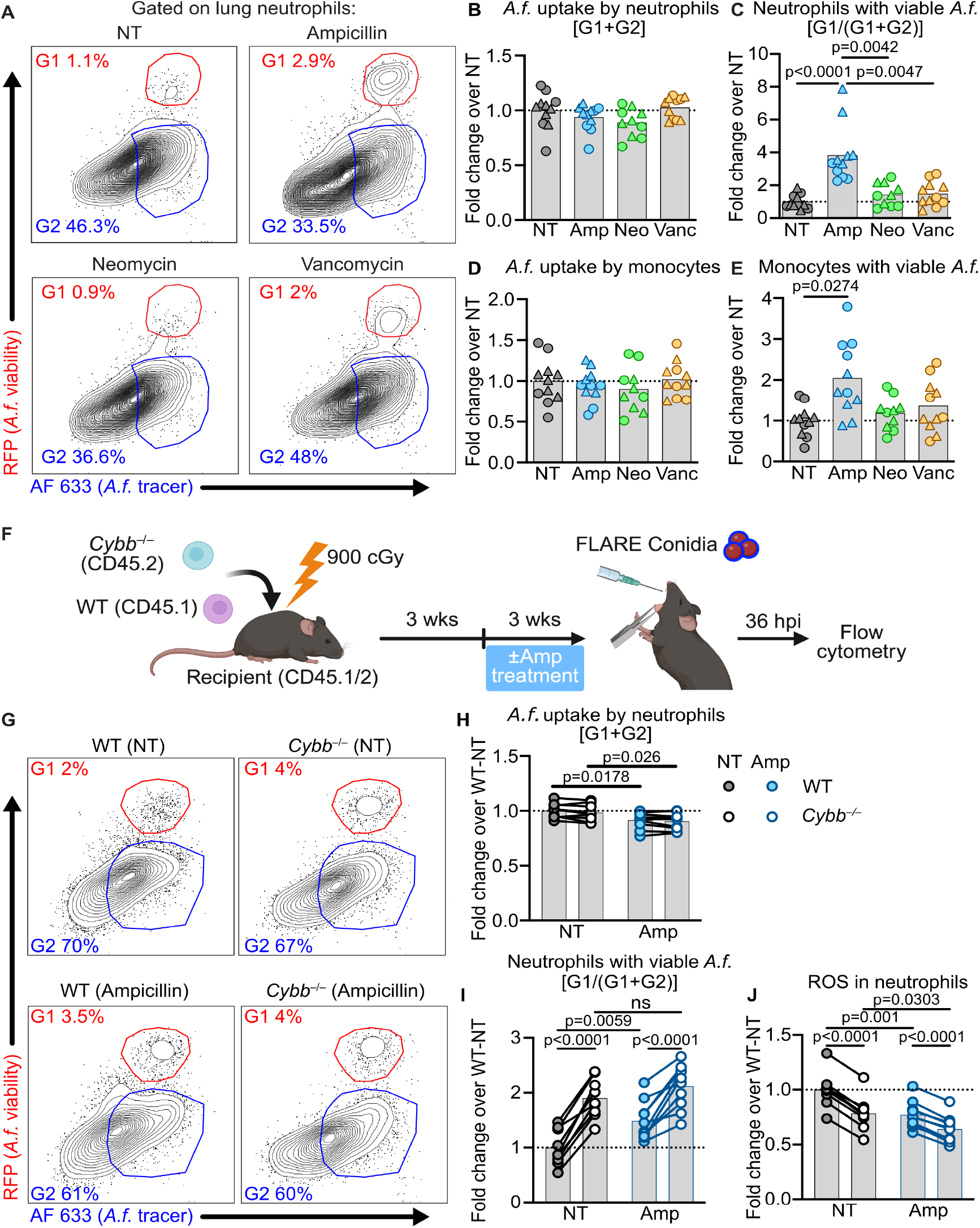
Ampicillin disrupts Nox2–dependent fungal killing by neutrophils. (A) Representative flow cytometry plots of lung neutrophils analyzed for RFP and AF633. (B and C) Quantification of *Aspergillus* uptake [G1 + G2] by lung neutrophils (Mean ± SD of NT: 55.67 ± 13.94) and lung monocytes (Mean ± SD of NT: 36.56 ± 11.15). (D and E) Quantification of lung neutrophils (Mean ± SD of NT: 2.33 ± 1.26) and lung monocytes (Mean ± SD of NT: 1.34 ± 0.89) with live *Aspergillus* viability [G1 / (G1 + G2)]. (F) Schematic of mixed bone marrow chimera generation followed by ampicillin treatment. Created with BioRender. (G) Representative flow cytometry plots of neutrophils analyzed for RFP and AF633 from chimeric mice. (H) Quantification of *Aspergillus* uptake [G1 + G2] by lung neutrophils from chimeric mice (Mean ± SD of NT-WT: 68.58 ± 4.65). (I) Quantification of *Aspergillus* viability [G1 / (G1 + G2)] in lung neutrophils from chimeric mice (Mean ± SD of NT-WT: 2.86 ± 0.93). (J) Quantification of ROS production by lung neutrophils from chimeric mice (Mean ± SD of NT-WT: 30810 ± 4841). Data in (B–E) and (H–J) are normalized to the average of the control group. Data in (B–E) are pooled from two experiments and the point shape indicates the experiment of origin. Data in (H– J) are from one experiment. Each point in (B–E) and (H–J) represents a mouse and the point shape in (B) represents the experiment of origin. Statistics: (B–E) Kruskal–Wallis test with Dunn’s multiple comparisons test. (H–J) RM two-way ANOVA, matched values are spread across a row and Uncorrected Fisher’s LSD, with a single pooled variance.

NADPH oxidase 2 (NOX2)-derived reactive oxygen species (ROS) is an essential mechanism used by human and murine neutrophils to kill *Aspergillus* (9, 10). Genetic defects in NOX2 result in greater susceptibility to *Aspergillus* infection in mice and humans (10, 11), underscoring the importance of this pathway to host defense. To test if ampicillin treatment impairs NOX2-dependent killing, we generated mixed bone marrow chimeras by reconstituting irradiated CD45.1^+^CD45.2^+^ mice with a 1:1 mix of CD45.1^+^ WT and CD45.2^+^ *Cybb*^−/–^ bone marrow cells. *Cybb* encodes the p91 subunit of NOX2. Mice were either left untreated or received ampicillin during the final 3 weeks of the 6-week reconstitution period and were challenged with 3×10^7^ FLARE conidia to assess fungal uptake and killing by WT and *Cybb*^*–/–*^ neutrophils in the same host.

In WT and *Cybb*^-/-^ lung neutrophils, we observed similar fungal uptake following no antibiotic treatment or ampicillin treatment (Figure 2G and Figure 2H). In contrast, fungal viability in lung neutrophils differed significantly with both genotype and antibiotic treatment. In untreated mice, a higher frequency of fungus-engaged *Cybb*^*–/–*^ neutrophils contained viable conidia compared to WT neutrophils (Figure 2G and 2I), consistent with published results (9). In ampicillin-treated mice, the frequency of WT neutrophils that contained viable *Aspergillus* conidia increased, mirroring the frequency observed in ampicillin-treated non-chimeric mice. In contrast, *Cybb*^*–/–*^ neutrophils did not show an increase in fungal viability with ampicillin treatment. Ampicillin thus impairs neutrophil fungal killing by disrupting NOX2-dependent fungal killing. Supporting this finding, ampicillin treatment significantly reduced global ROS production in WT lung neutrophils and, to a lesser extent, in *Cybb*^*–/–*^ neutrophils (Figure 2J).

Because ampicillin has been reported to directly scavenge ROS produced by neutrophils (12), we tested its direct effects on antifungal activity. Bone marrow cells were incubated with FLARE conidia and increasing concentrations of ampicillin spanning reported peak serum levels in rodents (13). Ampicillin treatment had no direct effect on neutrophil viability, neutrophil fungal uptake, or fungal killing by neutrophils (Figure S2), but treatment with the vacuolar ATPase inhibitor, bafilomycin A1, resulted in a significant increase in fungal viability in neutrophils, which is consistent with prior studies in macrophages (1) and confirmed that the assay could detect impaired fungal killing. Thus, ampicillin does not directly impair neutrophil antifungal activity.

Our findings contribute to a growing body of work demonstrating that antibiotic-induced dysbiosis of the microbiota impairs innate immune defense against pathogen challenge. Prior studies have shown that germ-free or broad-spectrum antibiotic-treated mice exhibit reduced neutrophil production and increased susceptibility to *Escherichia coli* and *Listeria monocytogenes* infection (14, 15); however, we observed no change in neutrophil accumulation in the lung following *Aspergillus* infection, suggesting that impaired cell abundance does not account for the ampicillin treatment-dependent susceptibility phenotype. Similarly, prior work found that neutrophils from antibiotic-treated neonatal mice retain normal phagocytic capacity, which we confirm in the context of *Aspergillus* (14). Instead, we identify a selective defect in fungal killing, consistent with a report that antibiotic treatment can impair neutrophil antimicrobial function against *S. pneumoniae* (16). Here, we identify reduced NOX2-dependent ROS production as a mechanism by which antibiotic treatment impairs neutrophil antimicrobial activity. Vancomycin treatment inhibited IL-17 production in the lung during *Aspergillus* infection (17). However, this effect was not associated with impaired fungal clearance or mortality, consistent with our finding that vancomycin does not alter fungal burden or survival. IL-17 levels in ampicillin-treated mice were not significantly different than untreated mice (Figure S3A). Antibiotic treatment reduced GM-CSF production in the lung during *S. pneumoniae* infection (4), which is notable because GM-CSF is essential for anti-*Aspergillus* immunity (18). However, GM-CSF levels were elevated, not diminished, in ampicillin-treated mice after *Aspergillus* infection (Figure S3B). This trend extended to other cytokines examined (Figure S3C–S3M), which were either comparable to or higher than in untreated controls. These data indicate that increased susceptibility arises from impaired phagocyte effector function rather than altered cytokine signaling.

Altogether, this study establishes a link between antibiotic treatment and increased susceptibility to *Aspergillus* infection, providing mechanistic insight into how antibiotic treatment impairs pulmonary antifungal immunity. These findings underscore the potential clinical relevance of antibiotic stewardship in immunocompromised individuals, who are at elevated risk for invasive aspergillosis, and suggest that selecting antibiotics that preserve microbiome integrity may mitigate infection-related morbidity and mortality in these high-risk settings.

## ACKNOWLEDGEMENTS

We thank Hohl Lab members for insightful discussions. These studies were supported by NIH grants F31 AI161996 (M.A.A.), P30 CA008748 (MSKCC, PI: Selwyn Vickers), R37 AI093808 (T.M.H.), and R01 R01AI191067 (T.M.H.). The funders had no role in the study design, data collection and analysis, decision to publish, or preparation of the manuscript.

## AUTHOR CONTRIBUTIONS

Mariano A. Aufiero, Conceptualization, Funding acquisition, Investigation, Writing original draft, Writing – review and editing. Tobias M. Hohl, Conceptualization, Funding acquisition, Supervision, Writing – original draft, Writing review and editing.

## DATA AVAILABILITY

Requests for further information, resources, or reagents should be directed to the corresponding author.

## SUPPLEMENTAL METHODS

### Mice

C57BL/6J wild-type (cat#: 000664) and *Cybb*^−/–^ mice (cat#: 002365) were obtained from The Jackson Laboratory (Bar Harbor, ME, USA). CD45.1^+^ mice (stock# 564) were purchased from Charles River Laboratories. CD45.1^+^CD45.2^+^ were generated by crossing CD45.1^+^ mice to C57BL/6J mice. Mice used in this study were 8-12 weeks old. Within experiments, mice were age- and sex-matched. Experiments were performed with both male and female mice. Mice were bred and housed in the Research Animal Resource Center at MSKCC in individual ventilated cages under specific pathogen–free conditions. Animal experiments were conducted with approval of the MSKCC Institutional Animal Care and Use Committee under the protocol 13-07-008.

### Antibiotic treatment

Antibiotics were administered to mice via drinking water for 3 weeks at the following concentrations: 1 g/L ampicillin, 1 g/L neomycin, or 0.5 g/L vancomycin as previously described (6). All antibiotics were purchased from the MSKCC pharmacy. Solutions were replaced weekly. The day prior to infection, mice were returned to normal drinking water.

### Mouse infection

*A. fumigatus* strain CEA10 (provided by R. Cramer, Dartmouth College) was used for all experiments. For experiments to analyze fungal uptake and killing using FLARE conidia, the CEA10–monomeric RFP strain was used. *A. fumigatus* conidia were grown on glucose minimal medium slants for 4–7 days at 37 °C before harvesting in phosphate-buffered saline (PBS) + 0.025% Tween 20 for experimental use. For FLARE experiments, 7.5 × 10^8^ conidia were incubated in EZ-Link Sulfo-NHS-LC-Biotin (10 μg/ml; Thermo Fisher Scientific) in 1 ml of 50 mM NaHCO3 buffer (pH 8.3) for 1–2 hours at room temperature, washed with 1 ml of Tris-HCl (pH 8) buffer, incubated with streptavidin (20 μg/ml) and Alexa Fluor 633 conjugate (Molecular Probes) in PBS for 45 minutes at room temperature, and resuspended in PBS + 0.025% Tween 20. Mice were anesthetized by isoflurane inhalation, and 3–6 × 10^7^ *A. fumigatus* conidia were instilled via the intratracheal route in 50 μl of PBS + 0.025% Tween 20.

### Quantification of lung fungal burden

To measure lung fungal burden, lungs were dissected and homogenized with a PowerGen 125 homogenizer (Thermo Fisher Scientific) for 10–15 seconds in 2 ml of PBS. A total of 10 μl was removed and diluted for plating onto Sabourand dextrose agar. Plates were incubated for 48 hours at 37 °C, and CFU were enumerated.

### Histology

Mouse lungs were collected into histology cassettes, stored in 4% paraformaldehyde for 16 hours, and then placed in ethanol. Samples were embedded in paraffin, sectioned, and stained with Grocott–Gömöri’s methenamine silver stain (GMS) to identify fungal organisms. Slides were scanned using a Pannoramic Digital Slide Scanner (3DHISTECH) using a 20×/0.8 numerical aperture objective.

### Flow cytometry

For analysis of immune cells, single-cell suspensions of mouse lungs were generated by putting lungs in gentleMACS C tubes and mechanically homogenizing in 5 ml of PBS using a gentle MACS Octo Dissociator (Miltenyi Biotec) in the absence of enzymes and then filtered through 100-μm filters. Red blood cells were lysed using red blood cell lysis buffer (Tonbo Biosciences), cells were blocked with anti-CD16/CD32, stained with fluorophore-conjugated antibodies, and analyzed on a Beckman Coulter Cytoflex LX. Single-color controls for compensation were generated using lung cells or OneComp eBeads Compensation Beads. Experiments were analyzed with FlowJo version 10.8.1. Dead cells were excluded with 4′,6-diamidino-2-phenylindole (DAPI) or eBioscience Fixable Viability Dye eFluor 506 (Thermo Fisher Scientific). The antibodies used are the following: anti-Ly6C (clone AL-21), anti-Ly6G (clone 1A8), anti-CD11b (clone M1/70), anti-CD11c (clone HL3), anti-CD45 (clone 30-F11), anti-CD45.1 (clone 104), anti-CD45.2 (clone A20), anti-I-A/I-E (clone M5/114.15.2), and anti-Siglec-F (clone E50-2440) all from Biolegend or BD Biosciences. Inflammatory monocytes were identified as CD45^+^ CD11b^+^ CD11c^−^ Siglec-F^−^ Ly6G^−^ Ly6C^hi^ MHC-II^−^ cells and neutrophils were identified as CD45^+^ CD11b^+^ Siglec-F^−^ Ly6G^+^. For experiments with mixed bone marrow chimeras, WT cells and *Cybb*^−/–^ cells were distinguished by CD45.1 and CD45.2 expression, respectively.

Conidial uptake and viability in lung phagocytes were assessed as previously described (8). After gating on lung monocytes or lung neutrophils, uptake was defined as the sum of RFP^+^AF633^+^ and RFP^−^AF633^+^ cells within each population. Conidial viability within a given leukocyte subset was calculated as the proportion of cells containing live conidia (RFP^+^AF633^+^) relative to all conidia-engaged cells (RFP^+^AF633^+^ and RFP^−^AF633^+^).

### Bone marrow chimeras

To generate mixed bone marrow chimeras, CD45.1^+^CD45.2^+^ recipient mice were irradiated (900 centigray) and reconstituted with a 1:1 mixture CD45.1^+^ wild-type and CD45.2^+^ *Cybb*^*–/–*^ bone marrow totaling 4 × 10^6^ donor bone marrow cells. After bone marrow transplantation, recipient mice were rested for 6 weeks prior to experimental use. Control mice received no antibiotics during this period, while ampicillin-treated mice were given antibiotics during the final 3 weeks, as described above.

### ROS Assay

Reactive oxygen species (ROS) production in lung neutrophils was measured using Dihydrorhodamine 123 (DHR; Invitrogen). Lung cells were incubated with 100 µM DHR, anti-Ly6G, and anti-CD11b antibodies in HBSS for 20 minutes at 37 °C. Dead cells were excluded with DAPI, and samples were analyzed on a Beckman Coulter Cytoflex LX.

### Ampicillin dose-response FLARE assay

Bone-marrow cells were harvested from the femurs and tibias of C57BL/6J mice, subjected to ACK lysis, and 1 × 10^6^ cells were seeded in a 24-well plate in complete RPMI-1640 (10 % heat-inactivated FBS, 2 mM L-gutamine, 1 mM sodium pyruvate, 10 mM HEPES, and non-essential amino acids. Pen/Strep was not added to the media for this experiment. Cells were pre-treated for 30 min at 37 °C with ampicillin (0, 1, 3, 10, or 30 µg/mL). Bafilomycin A1 (50 nM) served as an impaired-killing control. FLARE conidia were added at a MOI of 1. To prevent fungal germination, voriconazole (500 ng/mL) was added to the media. Following overnight culture, cells were labelled with anti-Ly6G, and anti-CD11b antibodies. Dead cells were excluded using DAPI. Samples were analyzed on a Beckman Coulter Cytoflex LX.

### Quantification of lung cytokines

Lungs were harvested and homogenized in 2 mL of PBS using a PowerGen 125 homogenizer (Thermo Fisher Scientific) for 10–15 seconds. Homogenates were centrifuged, filtered through a 40 µm cell strainer, and stored at –80 °C until analysis. Cytokine levels were measured using the LEGENDplex™ Mouse Inflammation Panel (13-plex) (BioLegend) according to the manufacturer’s instructions.

## SUPPLEMENTAL FIGURES

**Figure S1:**
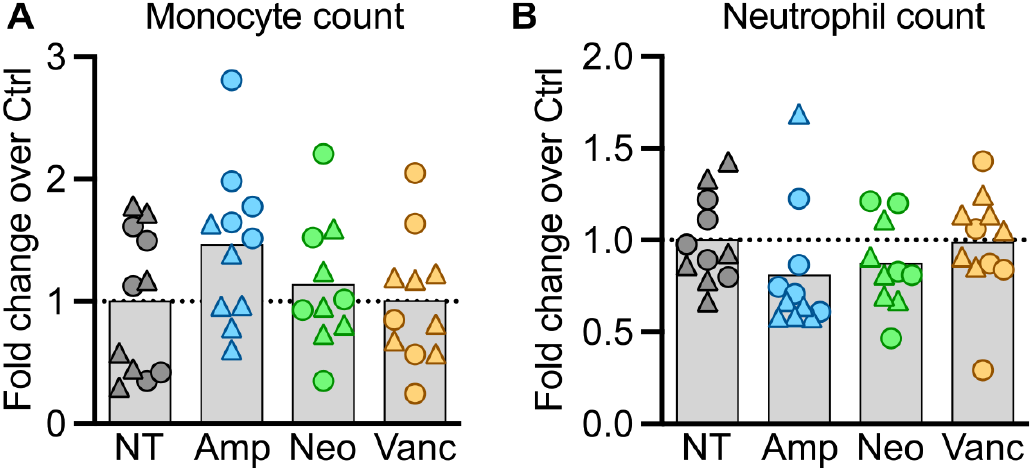
Quantification of lung phagocyte abundance and fungal uptake following *Aspergillus* infection. (A) Quantification of monocytes (Mean ± SD of NT: 104201 ± 66677) and (B) neutrophils (Mean ± SD of NT: 1632403 ± 444446) from the lungs of control or antibiotic-treated mice 36 hours after *Aspergillus* infection. Statistics: Kruskal–Wallis test with Dunn’s multiple comparisons test.

**Figure S2:**
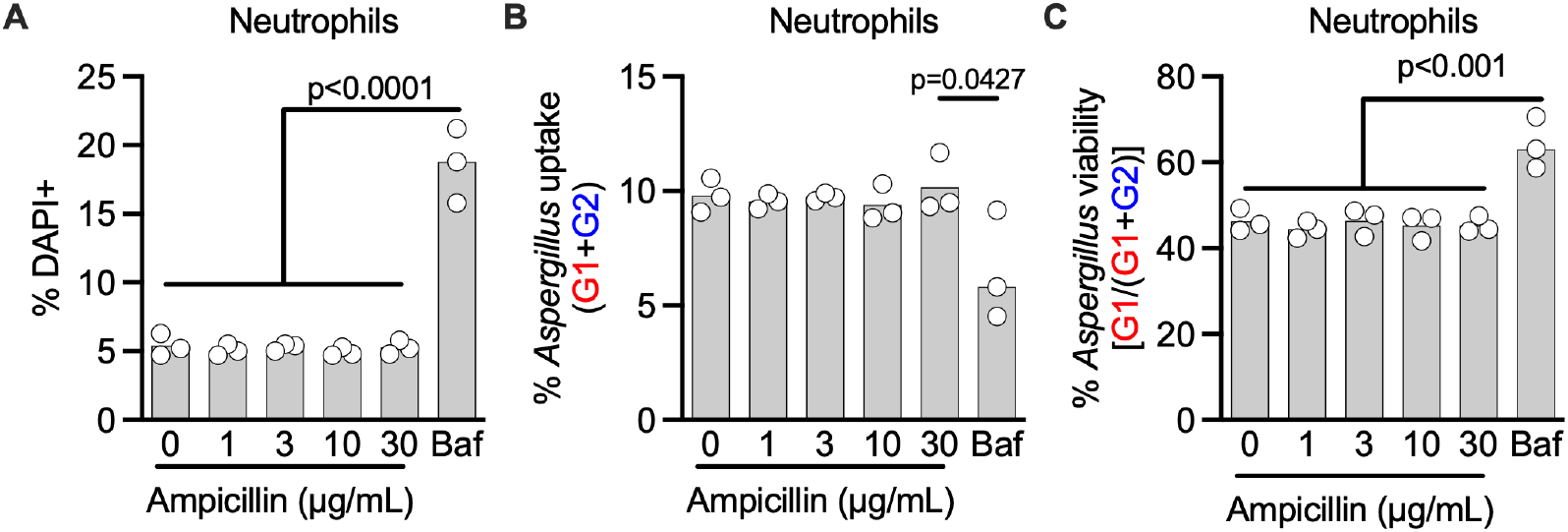
Ampicillin does not inhibit fungal killing by neutrophils in vitro. (A) Quantification of bone marrow neutrophil viability after overnight culture with FLARE conidia. (B) Quantification of *Aspergillus* uptake by DAPI^−^ bone marrow neutrophils. (C) Quantification of *Aspergillus* viability by DAPI^−^ bone marrow neutrophils. Data are representative of two independent experiments. Each point indicates a mouse. Statistics: Ordinary one-way ANOVA and Dunnett’s multiple comparisons test, with a single pooled variance.

**Figure S3:**
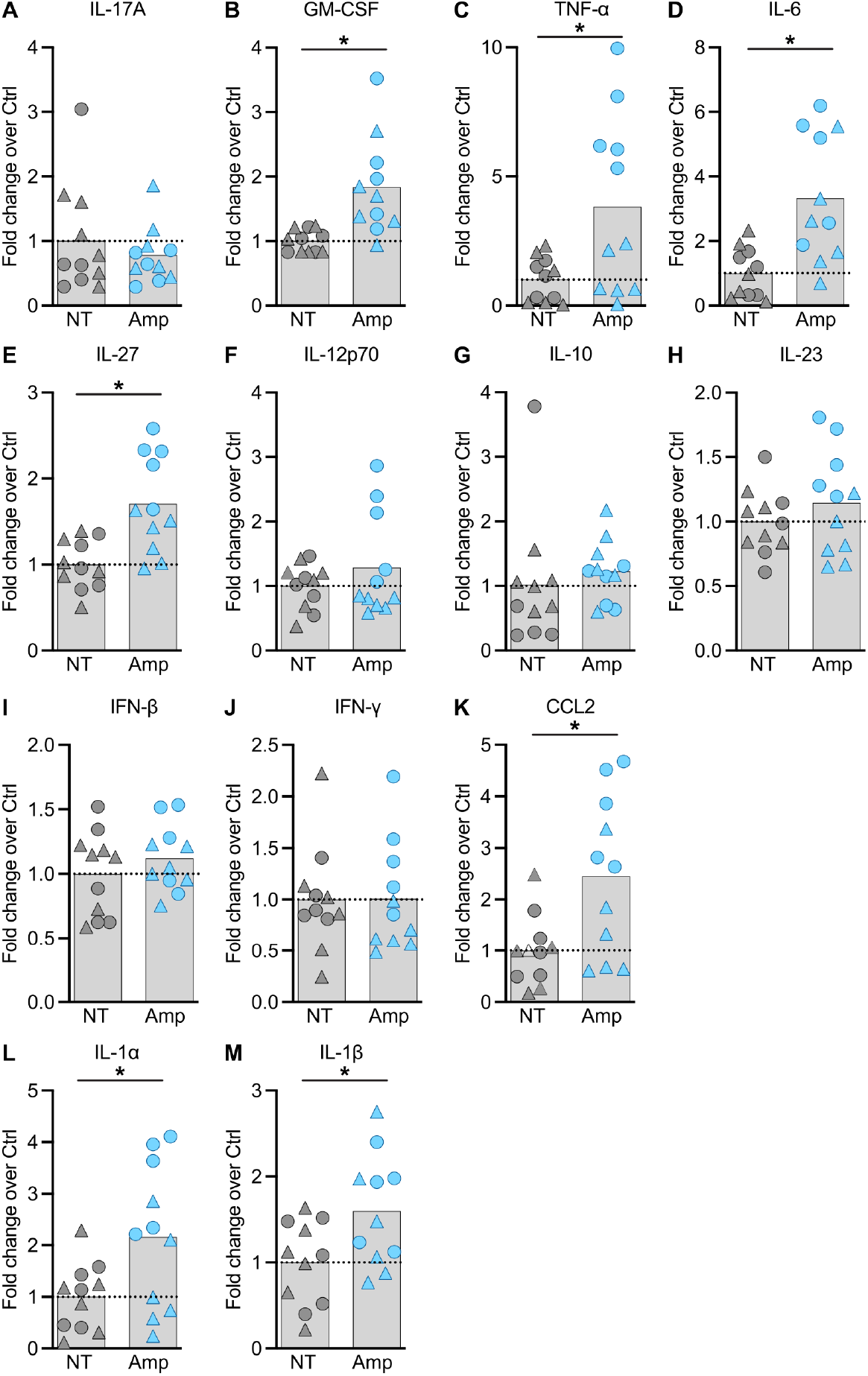
Lung cytokine levels in ampicillin-treated or control mice 36 hpi with *Aspergillus*. Data in (A-F) are normalized to the average of the control samples. Data are pooled from two experiments, and the point shape indicates the experiment of origin. Statistics: Unpaired t tests with Welch correction on each row, with multiple comparisons corrected using the Benjamini, Krieger, and Yekutieli procedure (FDR = 5%). Asterisks indicate differences that remained statistically significant after correction for multiple comparisons.

## Notes

### Competing Interest Statement

The authors have declared no competing interest.

